# Evaluation of a Polarization-Sensitive, Dual-Wavelength Wearable Photoplethysmography Sensor Across a Range of Skin Tones

**DOI:** 10.1101/2025.10.13.682143

**Authors:** Rutendo Jakachira, Wenyuan Yan, Sian C. Thomas, Yareli Macias-Sanchez, Lovisa Werner, Joshua A. Burrow, Shira Dunsiger, Kimani C. Toussaint

## Abstract

**Significance:** High-quality photoplethysmography (PPG) signals are essential for accurate extraction of cardiovascular metrics such as heart rate, heart rate variability, and perfusion index. However, signal degradation for individuals with dark skin tones can compromise PPG quality and pose challenges for equitable sensing.

**Aim:** We develop a dual-wavelength, polarization-sensitive PPG device to assess perfusion index (PI) across a range of skin tones.

**Approach:** To evaluate the impact of polarization on PPG signal quality, we record PI for co-polarized (polarized illumination and parallel-aligned polarized detection), and cross-polarized conditions (polarized illumination and orthogonally aligned polarized detection) at 655 nm and 940 nm in participants representing light, medium, and brown skin tone categories. Skin tone classification are based on the individual typology angle (ITA) values derived from the CIE L*b* color space measurements.

**Results:** At 940 nm, light from the cross-polarized light channel significantly increases PI (p < 0.05). At 655 nm, cross-polarization presents a statistically significantly enhanced PI (p < 0.05) relative to light from the co-polarized illumination condition, although the magnitude of the improvement decreases with lighter skin tone indication a possible interaction between skin tone and polarization. This improvement is consistent across all skin tones.

**Conclusions:** Our results suggest that the cross-polarized condition improves PPG signal quality by reducing the influence of superficial scattering and enhancing deeper vascular signals. This approach may be especially beneficial for individuals with darker skin tones and offers a promising path towards more robust and inclusive physiological monitoring using PPG-based technologies.

## Introduction

Photoplethysmography (PPG) is a non-invasive optical technique used to estimate arterial oxygen saturation and forms the foundation of all pulse oximetry measurements^1–4^. First introduced in the 1930s, PPG gained clinical utility in the 1970s with the Hewlett-Packard 47201A ear oximeter, the first device to continuously monitor oxygen saturation^5^. However, its complexity led to the development of the modern pulse oximeter by Dr. Takuo Aoyagi at Nihon Kohden in 1972, which became widely adopted by the mid-1970s^6^.

The underlying principle of PPG is based on the Beer-Lambert Law, which defines a linear relationship between light absorbance and the concentration of an absorbing molecule, assuming constant path length and wavelength^7^. When light interacts with the skin, absorption and scattering vary with the cardiac cycle: it increases during systole (due to greater blood volume and longer path length) and decreases during diastole^8^. The attenuated light that remains is detected, amplified, and processed to form the PPG waveform, which may include a dicrotic notch, a feature often visible in healthy, younger individuals with compliant arteries during the catacrotic phase^9,10^.

The waveform contains two main components: the DC signal, representing non-pulsatile tissues like chromophores within the tissue layers, muscle, veins, and bone, and the AC signal, reflecting pulsatile arterial blood. From this waveform, we derive the perfusion index (PI), the ratio of AC to DC components, which serves as a commonly used proxy for signal strength and vascular perfusion ^1,7,11–13^.

Today, pulse oximetry remains the primary clinical application of PPG. Since the 1990s, advances in signal processing and the integration of artificial intelligence and machine learning have enabled broader applications, such as sleep tracking, in consumer-grade wearables^14^. Thus, accurate PPG signals are essential for reliable oxygenation assessment in a variety of applications. However, growing evidence shows that pulse oximeters generally yield less accurate readings in individuals with dark skin tones, raising serious equity concerns in healthcare delivery^15,16^. While previous studies highlighting skin-tone related confounders in pulse oximeters have mainly focused on transmission-mode sensors^15^, and used race as a surrogate for pigmentation, our work builds on growing evidence that reflectance-mode sensors are not immune to this issue^17,18^. These challenges underscore the importance of device optimization to increase the accuracy of these devices ^14^. For example, adaptive filter methods such as the least square filter and adaptive step size least mean squares filter, are effective for real-time motion artifact removal to generally enhance signal-to-noise ratio (SNR) of the PPG signal. However, their removal heavily depends on accurate step size selection^19,20^. Additionally, wavelet-based methods such as discrete wavelet transform, have also been shown to enhance SNR at the cost of careful parameter tuning and increased computational complexity^21^.

A potential approach to improving the quality of the PPG signal could be the use of polarization gating. In the general context of tissue optics, it has been shown that for linearly polarized illumination, using co-polarized (parallel aligned) and cross-polarized (orthogonally aligned) detection permits separation (or gating) of surface reflection (superficial light) from multiple scattered (deep within the tissue) light, respectively^22^. This can be attributed to the fact that linearly polarized light propagating in skin undergoes multiple scattering with depth in the tissue which results in increased depolarization at depth. Using this approach, Lee et.al. demonstrated the performance of a flexible, wearable PPG sensor with the cross-polarized detection condition to reduce sensitivity to motion artifacts in the signal^23^. However, their device incorporated only a single wavelength, and the work did not explore the response across different skin tones nor was the PI determined. Increasing the source-detector separation can also increase sampling depth, and this is indeed a widely used approach. However, there is a tradeoff between the penetration depth and signal quality as when the source and detector separation are increased beyond a certain point^24^. Cross-polarization provides a compact, filter-based alternative that achieves some depth discrimination without increasing device size or an important consideration in wearable or miniaturized systems.

In this work, to quantify the impact of polarization on PPG signal quality, we develop a flexible, wireless, dual-wavelength, wearable PPG to measure the PI across various skin tones, using co-polarized (polarized illumination and parallel-aligned polarized detection), and cross-polarized conditions (polarized illumination and orthogonally aligned polarized detection) at 655 nm and 940 nm across a range of skin tones. To assess skin tone, we employ the individual typology angle (ITA)^25,26^. We find generally that the cross-polarization condition yields a statistically significant higher PI than the co-polarized condition (p < 0.05). Our preliminary findings support the incorporation of polarized illumination and detection as a means of improving the quality of the PI, which can inform the development of more accurate pulse oximeters for dark skin tones.

## Methods

### Device Design

Figure 1 illustrates the design and deployment of the wearable, polarization-enhanced PPG device. As shown in Fig. 1a, the top silicone substrate contains rectangular cutouts that align with the underlying polarization layer, which separates reflected light into co- and cross-polarized channels before reaching the photodiodes (PDs). The fPCB is fabricated from a 100-µm thick polyimide interlayer and flanked by two 18-µm thick patterned copper layers. The board hosts several key components, including: a tri-wavelength LED package (SFH 7016, OSRAM), with the 540-nm wavelength channel unused; four silicon PIN photodiodes (VEMD 5080, Vishay Semiconductors); a Bluetooth Low Energy (BLE) system-on-chip (SoC) module (ISP1807, Insight SIP); a power management integrated circuit (nPM1100, Nordic Semiconductor); an optical analog front-end (MAX86141, Analog Devices); a tuned copper coil; and associated passive components. To address potential concerns about optical crosstalk, we implement a time-multiplexed LED activation scheme, ensuring that only one LED was active at any given time. A 12 µs settling time is introduced between LED turn-on and ADC integration to allow the LED output to stabilize, and each photodiode readout was captured during a dedicated 58.7 µs integration window. The flexible board folds into a compact configuration suitable for wrist-worn operation, as depicted in Fig. 1b. The optical interaction between the device and biological tissue is illustrated in Fig. 1c. Light emitted by the LEDs is first linearly polarized by the polarization film and then propagates through the epidermal and dermal layers of the skin. As the light travels through tissue, it undergoes scattering and absorption by surrounding tissue, vasculature and chromophores such as melanin, and hemoglobin. This configuration enables concurrent acquisition of both co- and cross-polarized channels.

**Figure 1:**
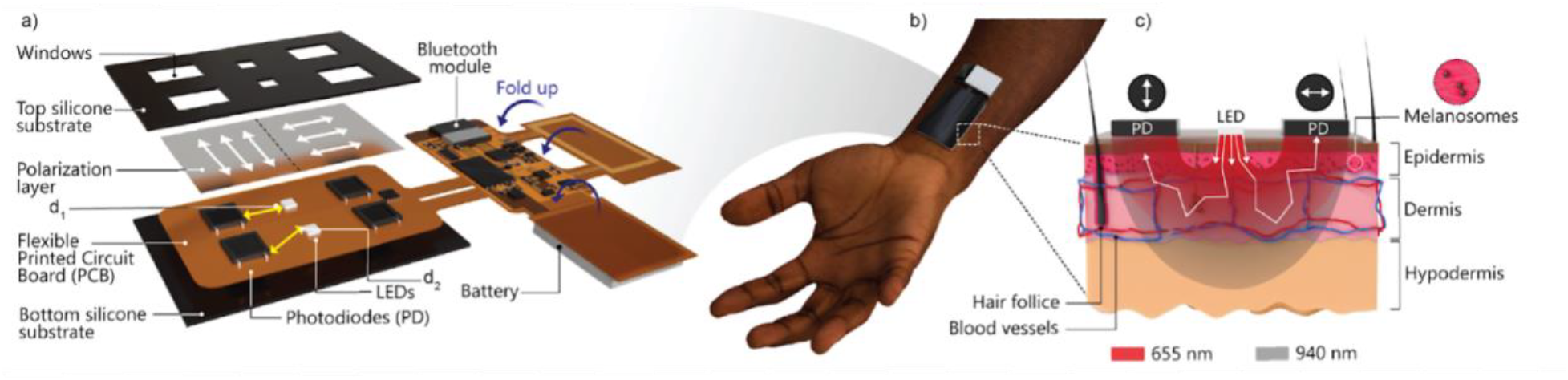
Design and application of a polarization-sensitive wearable device for PI measurements across skin tones. a) Exploded schematic of the device architecture, comprising a multilayer stack with top and bottom silicone substrates, a polarization layer, fPCB, LEDs, and PDs. The windowed top layer aligns with optoelectronic components to allow light transmission. Two source-detector separations are used: d_1_= 9 mm and d_2_ = 10.82 mm. An image taken from the actual study is displayed in Fig. S1. b) Image of the device worn on the forearm, demonstrating skin contact and flexibility. Adjacent blocks of the fPCB are folded during packaging to reduce the overall size of the device (6.2 cm x 2.4 cm x 0.8 cm). c) Schematic cross-section of skin showing light propagation from LEDs (655 nm and 940 nm) through the epidermis, dermis, and hypodermis, with differential absorption and scattering by melanosomes and vasculature. Photodiodes positioned with polarizers, enabling capture of co- and cross-polarized signals to isolate superficial versus deeper tissue contributions, respectively.

### Electronics architecture

The electronic components of the wearable system are distributed across a fPCB, which coordinates optical emission, signal acquisition, onboard processing, and wireless communication, as illustrated in Fig. 2. The device LEDs are controlled by a programmable LED driver. Four silicon photodiodes (PD1–PD4) are symmetrically positioned on either side of the LED module, and these photodiodes are interfaced with transimpedance amplifiers (TIAs), which convert photocurrent10/13/2025 11:07:00 PM into voltage signals. The resulting analog signals are digitized using high-resolution, 19-bit charge-integrating analog-to-digital converters (ADCs) operating at 25 samples per second and then passed to the onboard microcontroller for processing. During operation, the LEDs are driven at 14.53 mA. The entire system is powered by a 60 mAh lithium-polymer battery, which supports continuous operation for up to 24 hours

**Figure 2:**
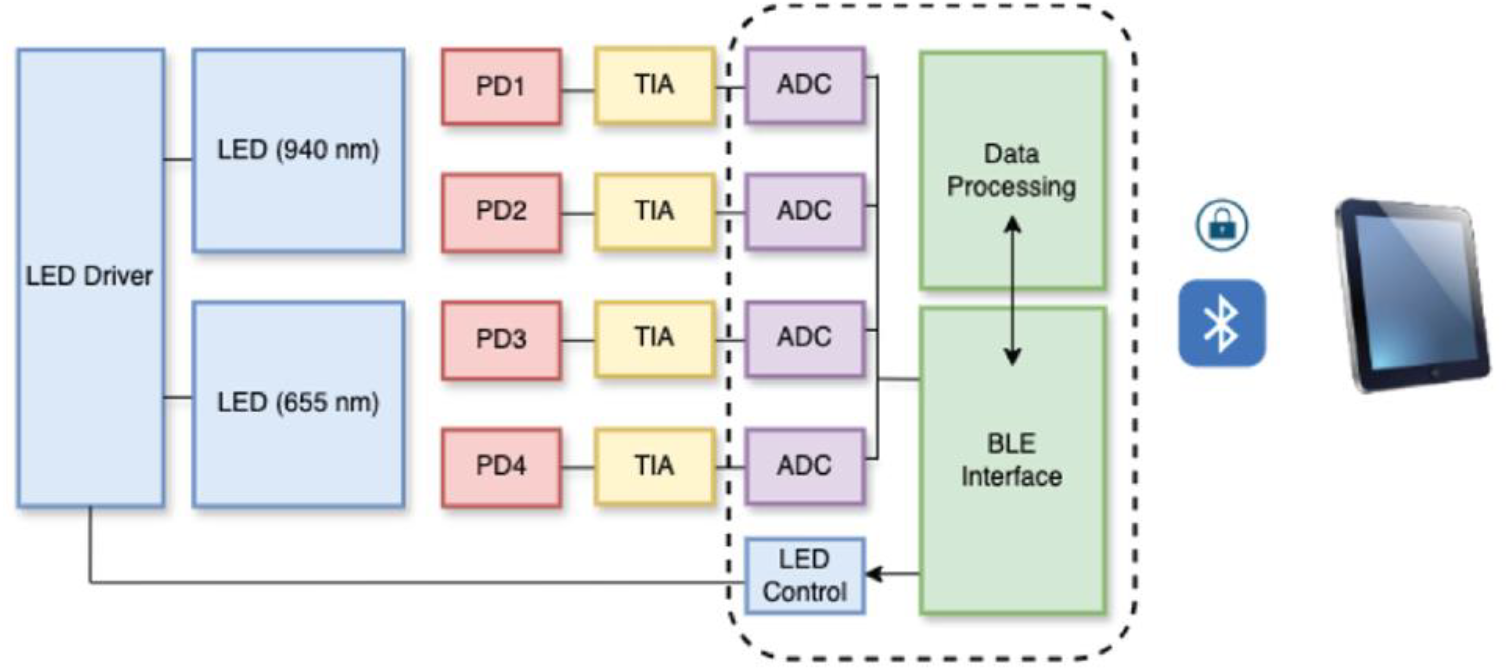
System architecture of the wearable PPG device.

### Algorithm Structure

The acquired raw PPG signal undergoes a multi-stage processing pipeline to extract mean PI values from each volunteer, as shown in Fig. 3. The raw data is exported from an iPad to a PC, where it is analyzed using MATLAB. The signal is first subjected to low-pass filtering using MATLAB’s built-in lowpass () function, with a cutoff frequency of 5 Hz. This function applies a zero-phase, finite impulse response filter that attenuates frequency components above 5 Hz while preserving key physiological features, such as the cardiac pulse, which typically resides in the 1– 2 Hz range. The -3 dB cutoff ensures that frequency components at 5 Hz are reduced to approximately 70.7% of their original amplitude, effectively minimizing high-frequency noise while maintaining waveform morphology.

**Figure 3:**
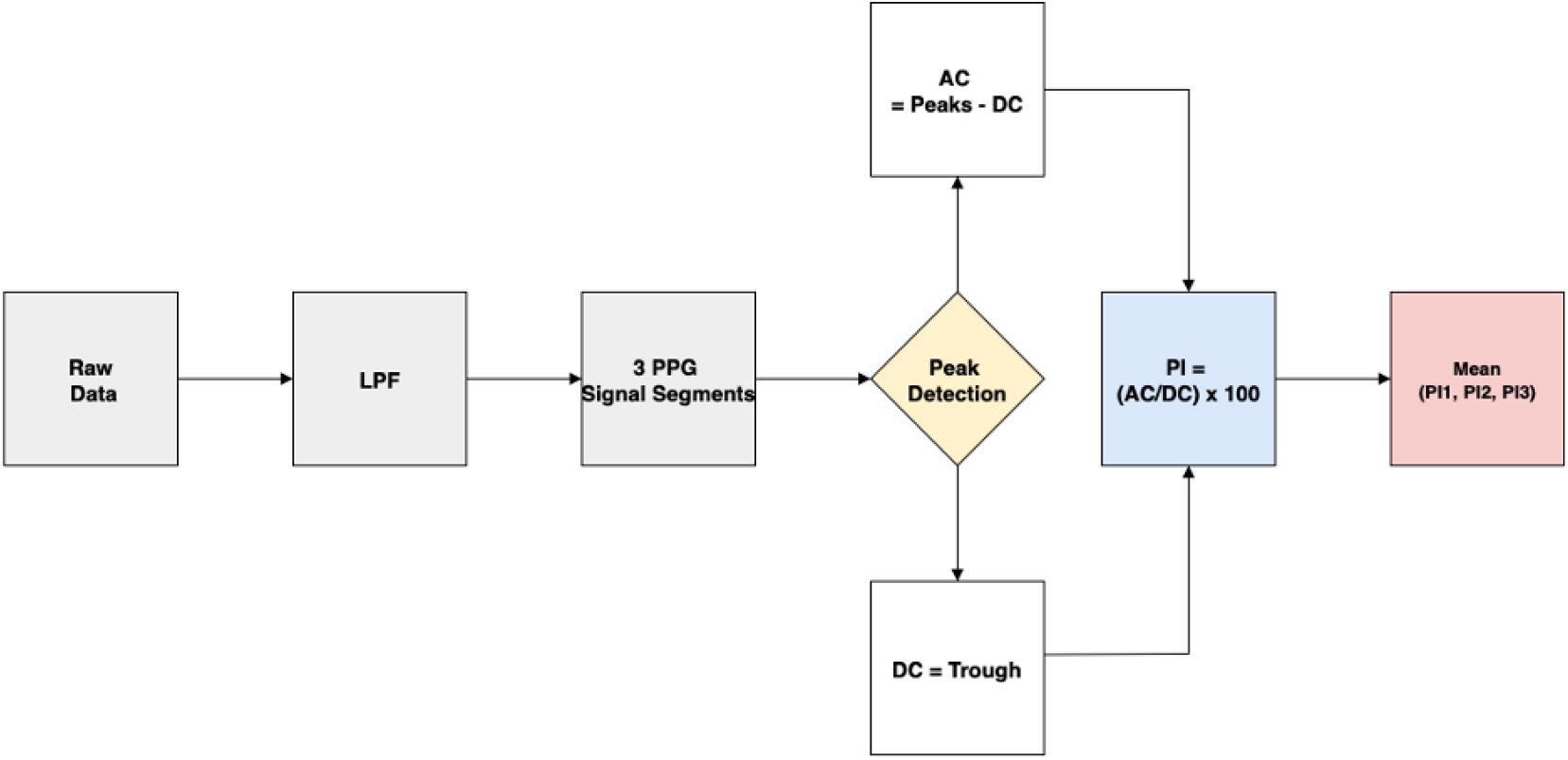
Signal processing system diagram for PI computation from PPG data.

To improve signal quality in the presence of involuntary motion artifacts, the sampling rate parameter in the lowpass () functions is dynamically adjusted between 20 and 50 Hz depending on the quality of the raw signal, to ensure the greatest signal fidelity. This helps optimize the performance of the peak-finding algorithm by balancing time resolution and noise suppression. To ensure detected peaks are true peaks, we assess the frequency at which peaks occur, keeping in mind that the typical heart rate frequency for an adult aged 18-30 years is 60-100 beats per minute ^27^. After filtering, the signal is segmented into three 10-second PPG waveforms to ensure consistent identification of peaks and troughs for downstream PI calculations.

Peak detection is performed on each segment using the findpeaks() function in MATLAB. This allows the extraction of the pulsatile (AC) and baseline (DC) components of the PPG signal. The DC value is defined as the local trough (minimum), while the AC value is computed as the difference between the identified peak and the preceding trough (DC) such that *AC* = *Peaks* – *DC*. The PI values for each segment are calculated as the ratio of the AC value and corresponding DC value expressed as a percentage. Consequently, the mean PI value is then obtained by averaging the PI values for the three segments, P_1_, P_2_, and P_3_. This process is repeated for the co-, and cross-polarized channels.

### Recruitment protocol

This Institutional Review Board (IRB) at Brown University granted a waiver of IRB approval for this study, as it focuses solely on device calibration and does not meet the federal definition of research intended for generalizable knowledge. A total of nine healthy volunteers representing a range of skin tones were recruited and categorized into three groups: light, medium, and brown. Skin tone classification was performed using a Colorimeter DSM-4 which measures skin reflectance and computes values based on the CIELAB color space. Specifically, the ITA is derived from the L* and b* (components of the CIE L*a*b* color space and is widely used in dermatological studies to quantitatively classify skin tone. In this study, ITA values above 23° indicate light skin, values between –5° and 23° denote medium skin tones, and values below –5° correspond to brown skin tones. Like the approach taken by Leeb et. al, we apply alternative ITA cutoffs to those traditionally used to address that fact that we have a limited number of participants below –30°^28^.

### Data acquisition protocol

Upon arrival, the volunteers are seated in a designated common area within the study location and given five minutes of rest to allow their heart rates to stabilize. Skin-tone measurements are then taken at the wrist, specifically at the site of PPG device placement. For each volunteer, 20 consecutive ITA measurements are recorded, and the mean ITA value is used for analysis.

Following the skin-tone assessment, the PPG device is placed on the left wrist. To ensure firm adhesion at the skin-sensor interface, the device is attached to volunteers’ wrist using medical-grade transparent film dressing (Tegaderm, 3M). Measurements are initiated with 5 minutes of baseline recording. After the readings are stabilized, volunteers are asked to remain still. Two minutes of continuous PPG data is then collected. This allows for a direct comparison of mean PI values between co-, and cross-polarized conditions across different skin tones.

### Statistical Analysis

To analyze the impact of polarization condition and skin tone on PI, we used a series of linear regression models implemented with generalized estimating equations (GEE) with robust standard errors. GEEs are a statistical method well-suited for handling correlated or clustered data, such as repeated measures within individuals. This approach accounts for the within-subject correlation that arises when participants contribute multiple observations and adjusts standard errors to reflect the non-independence of observations within participant. Models regressed PI outcomes on ITA, polarization condition and ITA*condition. All analyses were conducted using R (RStudio Team, 2020), with significance level set at 0.05 a priori.

## Results

Figure 4 displays the ITA values at the wrist for study participants, plotted in the L*b* plane of the CIE 1976 (L*, a*, b*) color space. Each point represents the average skin color measurement for a participant, with error bars indicating variability in L* across 20 repeated scans. ITA values, derived from L* and b* measurements, are used to classify skin tone independent of race or ethnicity. Participants with lighter skin tones exhibit more positive ITA values, while those with darker skin tones have more negative ITAs.

**Figure 4:**
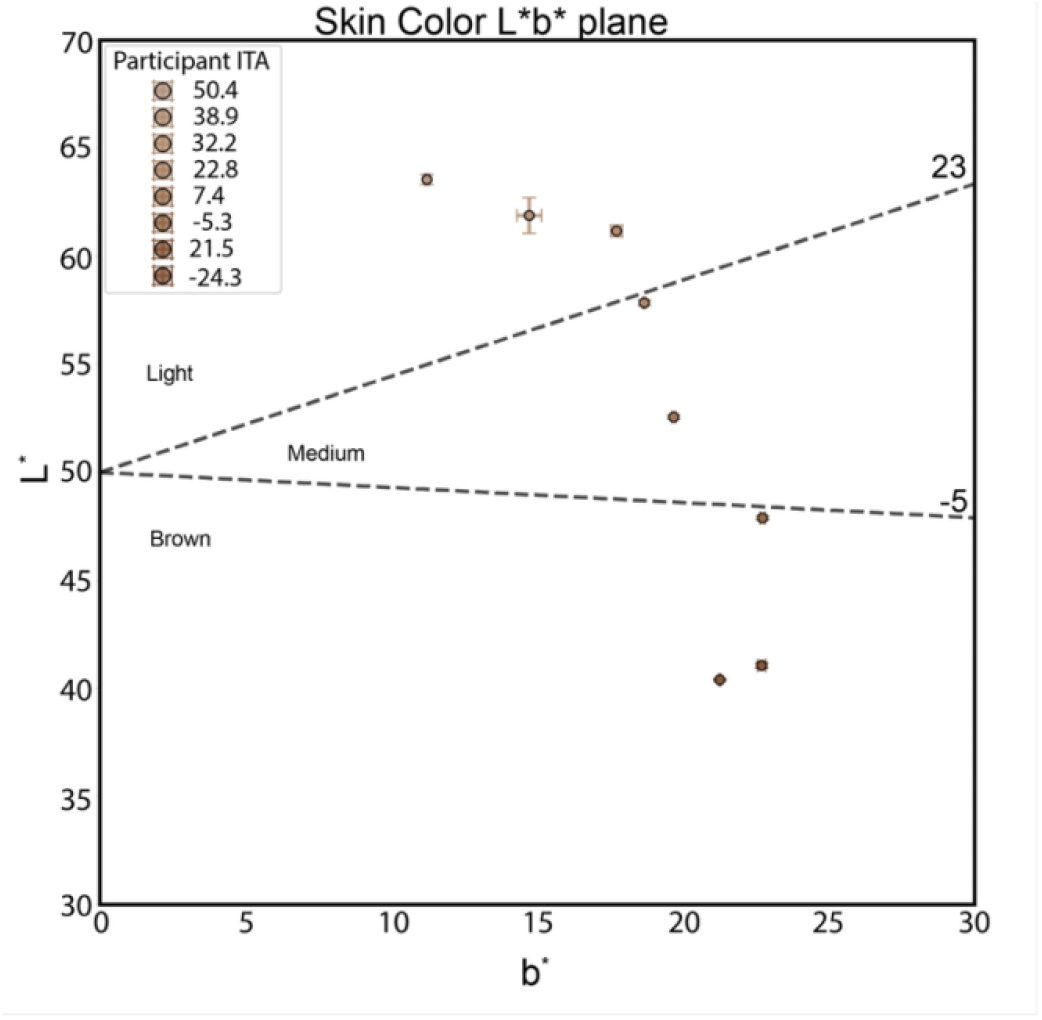
Participant skin tone distributed in the CIE L*b* color space.

The study initially included nine participants; however, one participant was excluded due to having a substantially higher perfusion index compared to the rest of the cohort. This is because the participant had engaged in strenuous physical activity shortly prior to participating in our study. This was not true for the other participants and thus we felt confident with excluding this individual from the analysis to enable a fairer comparison across subjects under similar physiological conditions. For the remaining 8 participants, the mean and standard deviation ITA values are summarized in Table 1.

**Table 1:** Participant ITA Summary.

| Participant # | ITA (°) | Standard Deviation (°) |
| --- | --- | --- |
| 1 | 50.4 | 0.4 |
| 2 | 38.9 | 1.5 |
| 3 | 32.2 | 0.6 |
| 4 | 22.8 | 0.3 |
| 5 | 7.4 | 0.2 |
| 6 | -5.3 | 0.2 |
| 7 | -21.3 | 0.4 |
| 8 | -24.3 | 0.1 |

In Fig. 4, the black dashed lines at 23° and –5° represent ITA thresholds for classifying skin tone: values above 23° correspond to “light” skin tones, between –5° and 23° to “medium,” and below –5° to “brown.” Based on this classification, three participants fell within the light category, two in the medium, and three in the brown category. This spread highlights the inclusion of a range of skin tones in the study and supports the use of ITA as an objective metric for skin tone classification in optical device validation.

Figure 5 shows normalized PPG signals over 10-second windows for three representative participants spanning the light, medium, and brown skin tone categories (panels a–c, respectively). Signals are shown across four measurement conditions, combining two polarization conditions: cross-polarized, and co-polarized, with two wavelengths: 655 nm and 940 nm. Each stacked plot corresponds to one of these four combinations. The left y-axis displays PPG signals normalized within each subplot to highlight waveform morphology, while the right y-axis shows the corresponding unnormalized intensity in arbitrary units (au), enabling direct comparison of signal amplitude across conditions. It should be noted that we plot detected light intensity as a function of time without inversion. In reflectance PPG, systole increases arterial blood volume and absorption, reducing detected intensity; during diastole, absorption decreases and detected intensity increases^29^. For the light-skinned participant, unnormalized signal ranges are approximately 45,900–46,400 au (655 nm, co-polarized), 45,120–45,640 au (940 nm, co-polarized), 38,080–38,920 au (655 nm, cross-polarized), 30,210–30,780 au (940 nm, cross-polarized).

**Figure 5:**
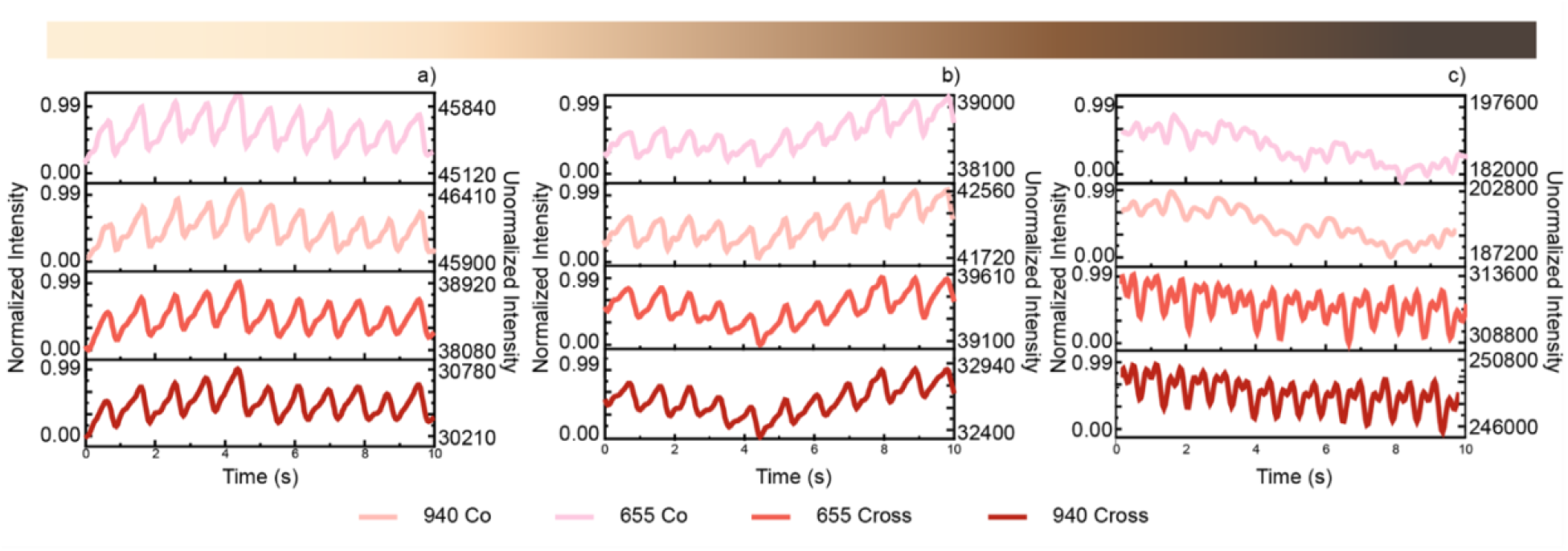
PPG waveforms from three participants in the light, medium and brown skin tone categories (a-c), respectively.

These three participants selected represent a range of ITA-based skin-tone categories in our study. Data from the remaining five participants are provided in Supplementary Figure S2, which includes the remaining set of PPG traces across all four optical conditions.

The bar graphs in Fig. 6 show the mean PI values measured using cross-polarized, and co-polarized conditions at 655 nm (left) and 940 nm (right). Across both wavelengths, cross-polarized measurements consistently yield higher PIs. This suggests improved signal recovery under cross-polarized conditions, potentially mitigating the impact of increased melanin content on PPG signal quality. This is confirmed by statistical analysis performed at these wavelengths as shown in Tables 2 and 3.

**Table 2.** 655 nm GEE model output.

| Coefficients | Estimate | Standard error | 95% CI | Wald | Pr(>\|W\|) |
| --- | --- | --- | --- | --- | --- |
| Intercept | 0.66246 | 0.10416 | 0.409, 0.916 | 40.45 | 2*10<sup>-10</sup> *** |
| ITA | 0.00584 | 0.00281 | -0.001, 0.013 | 4.33 | 0.0374 |
| Cross-polarized | 0.82845 | 0.26958 | 0.226, 1.431 | 9.44 | 0.0021 ** |
| ITA*Cross-polarized | -0.01531 | 0.00968 | -0.035, 0.004 | 2.50 | 0.1139 |

**Table 3.**
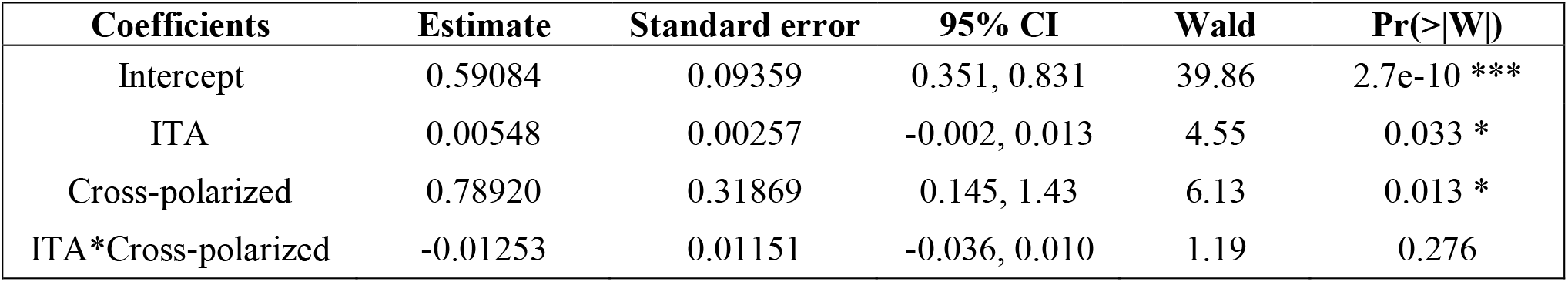
940 nm GEE model output.

| Coefficients | Estimate | Standard error | 95% CI | Wald | Pr(>\|W\|) |
| --- | --- | --- | --- | --- | --- |
| Intercept | 0.59084 | 0.09359 | 0.351, 0.831 | 39.86 | 2.7e-10 *** |
| ITA | 0.00548 | 0.00257 | -0.002, 0.013 | 4.55 | 0.033 * |
| Cross-polarized | 0.78920 | 0.31869 | 0.145, 1.43 | 6.13 | 0.013 * |
| ITA*Cross-polarized | -0.01253 | 0.01151 | -0.036, 0.010 | 1.19 | 0.276 |

**Figure 6:**
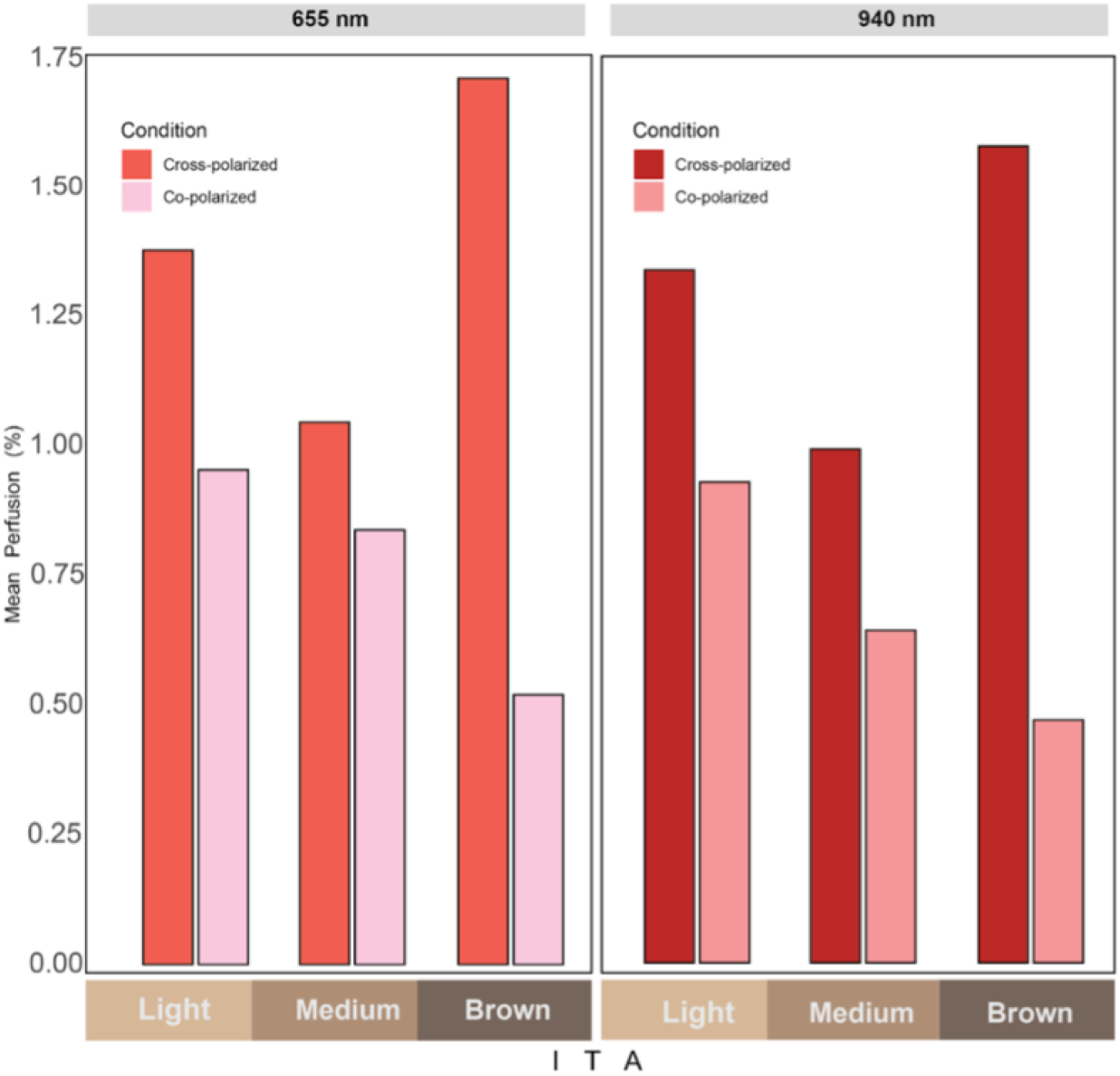
Mean PI across skin tone categories for each polarization condition and wavelength.

Models estimates at 655 nanometers shown in table 2 indicate a significant difference between cross-polarized and co-polarized conditions (p<0.05) such that the mean perfusion index is significantly higher for the cross-polarized condition compared to the co-polarized condition. Although not statistically significant, there is a trend to suggest that the difference between the mean perfusion indices for cross-polarized and co-polarized might vary by skin tone. These effects should be further tested in a larger sample as the current sample is under-powered to formally test statistical moderation, but the point estimates provide some evidence in support of further examination. The point estimates suggest that the lighter the skin tone is, the less of a difference there is between the cross-polarized and co-polarized conditions.

The estimates from our Generalized Estimating Equations (GEE) model at 940 nm shown in table 3 show that the cross-polarized condition has a larger perfusion index in relation to co-polarized condition (p<0.05). This is a similar trend to what we observed at 655 nm. Results do not indicate a significant conditional effect of skin tone (p=.276). However, point estimates are in the direction the same direction as the 655 nanometers wavelength. This effect should be further tested in a fully powered sample.

## Discussion

Our results suggest that the cross-polarization condition effectively reduces superficial scattering and enhances signal contributions from deeper pulsatile vasculature, leading to improved SNR. This is encouraging as it indicates the potential to enhance SNR in signals confounded by higher concentrations of melanin in the skin. By improving signal, the polarization-sensitive designs inform an approach to mitigate the racial and ethnic disparities in PPG-based technologies such as pulse oximeters, heart rate monitors and sleep apnea diagnostic tools. Given the pilot nature of this study, these findings are very preliminary but highlight meaningful trends that warrant further investigation with a larger and more diverse cohort. A limitation of our current approach is the unintended consequences of cross polarization. While the use of cross-polarization may lead to a greater focus on the deeper vessels rather than superficial epidermis where melanin resides, nonpulsatile venous blood is deeper and could also serve as a factor that makes pulse oximeters less accurate. Future work will aim to characterize and account for these deeper-tissue contributions more explicitly.

## Conclusion

In this study, we created a flexible, wireless, dual-wavelength, PPG wearable and used it to demonstrate that polarization gating, particularly employing cross-polarized illumination and detection, provides a statistically significant improvement of the quality of PPG signals as measured by the PI, across a range skin tone.

These results highlight the potential of polarization-based optical techniques to reduce bias in clinical PPG-based devices, particularly for individuals with darker skin tones who have historically been underserved by conventional pulse oximetry. Future work includes expanding our work to include more participants, exploring measurements at different anatomical sites, and incorporating the unpolarized condition on the same wearable as the polarized conditions. Furthermore, the use of polarization vector beams could offer polarization diversity in a single shot, as we have previously shown^30^, which will permit exploration of illuminating with multiple states of polarization in parallel and offers a path for polarization optimization for PPG detection.

## Supporting information

Figure S1. The device deployed on the wrist and held in place using clinical adhesive tape to ensure stable contact during data collection.

Figure S2. Remaining set of PPG traces across 6 optical conditions from light to brown(left to right).

## Appendix A: Supplemental Material

## Code and Data Availability

The datasets supporting the conclusions of the article are available on GitHub (https://github.com/10dojak/PPG_data). Additional information may be obtained from the authors upon reasonable request.

## Acknowledgements

R. J. acknowledges support from the Google PhD fellowship program in Health Research and Algorithmic Fairness. J.A.B. acknowledges support from the Burroughs Wellcome Fund Postdoctoral Enrichment Program (Grant No. 009248). Additionally, this project has been made possible in part by a grant from the Chan Zuckerberg Initiative DAF, an advised fund of Silicon Valley Community Foundation. We also acknowledge partial support from the Brown University Center for Digital Health. Finally, we gratefully acknowledge Jacob Trueb and John Rogers at Northwestern University for valuable assistance with the data acquisition software. We are grateful to Lydia Vignale for her advice regarding the participant data-taking protocol

**Rutendo Jakachira** received her B.S. degree in physics from Drew University in 2019 and is currently working towards her Ph.D. in physics at Brown University working in the PROBE Lab. Her current research interest lie in biomedical opto-electronic devices, signal processing and MonteCarlo modeling of turbid media.

**Wenyuan Yan** received his B.S. in Materials Science and Engineering from Tongji University (2020) and an M.S. from Northwestern University, where he worked on wearable bioelectronics for oxygenation monitoring. He is currently at Cornell University pursuing a Ph.D. in Materials Science.

**Sian C. Thomas** has a degree in biomedical engineering and computer science from Brown University. Her research focuses on improving the functionality of wearable devices and developing signal processing algorithms for SpO_2_ readings.

**Yareli Macias-Sanchez** received her Sc.B. degree in Biomedical Engineering from Brown University in 2025. She is currently working in the PROBE Lab at Brown University on the development of a pulse oximeter that provides accurate measurements across different skin tones. Her research interests include biomedical optics, medical device design, and health equity in diagnostic technologies.

**Lovisa Werner** is a Biostatistician within the Survey, Qualitative and Applied Data Research Core at Brown University School of Public Health. She received her B.A. degree in biology from Connecticut College in 2022 and her Master of Public Health in epidemiology from Brown University in 2024. She works with data gathered via medical and behavioral research, to formulate conclusions and make predictions.

**Joshua A. Burrow** earned dual B.S. degrees in amthematics and physics from Morehouse College (2017) and Ph.D (2021) in Electro-Optics & Photonics from the university of Dayton. At Brown University, he was a Burroughs Wellcome Fund Fellow in the Probe Lab. His research focused on light-matter interactions, nanophotonics, meta-optics, polarization microscopy, and biophotonics.

**Shira Dunsiger** is biostatistician with significant experience analyzing data from behavioral medicine, including physical activity, sleep, smoking, mood, violence, adherence, weight loss, alcohol and nutrition. Her interests are in identifying statistical methods that address the challenges in this type of data. Shira has significant experience working with high dimensional longitudinal data, including EMA, wearables and intensive sampling outcomes.

**Kimani C. Toussaint, Jr**. is the Thomas J. Watson, Sr. Professor and Senior Associate Dean in in Brown’s School of Engineering and Director for the Center for Digital Health. He leads the PROBE Lab, an interdisciplinary group advancing nonlinear optical imaging for biological tissue assessment and developing plasmonic nanostructures for light-driven control of matter.

