## Supplementary figures and images for "Evaluation of a Polarization-Sensitive, Dual-Wavelength Wearable Photoplethysmography Sensor Across a Range of Skin Tones"

### Figure S1. The device deployed on the wrist and held in place using clinical adhesive tape to ensure stable contact during data collection.

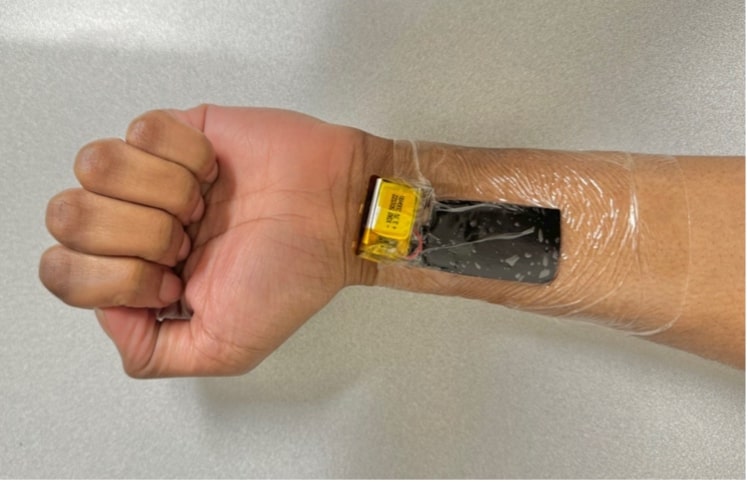

### Figure S2. Remaining set of PPG traces across 6 optical conditions from light to brown(left to right).

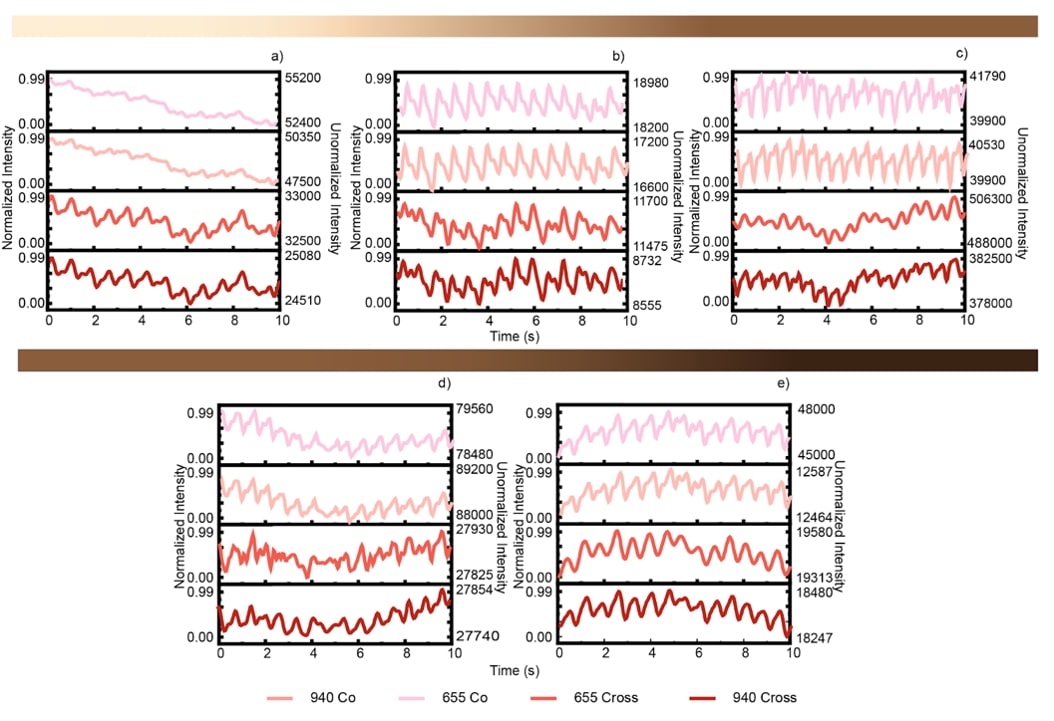
